# Enhanced Behavioral Performance through Interareal Gamma and Beta Synchronization

**DOI:** 10.1101/2023.03.06.531093

**Authors:** Mohsen Parto-Dezfouli, Julien Vezoli, Conrado Arturo Bosman, Pascal Fries

## Abstract

Cognitive functioning requires coordination between brain areas. Between visual areas, feedforward gamma synchronization improves behavioral performance. Here, we investigate whether similar principles hold across brain regions and frequency bands, using simultaneous local field potential recordings from 15 areas during performance of a selective attention task. Short behavioral reaction times (RTs), an index of efficient interareal communication, occurred when occipital areas V1, V2, V4, DP showed gamma synchronization, and fronto-central areas S1, 5, F1, F2, F4 showed beta synchronization. For both area clusters and corresponding frequency bands, deviations from the typically observed phase relations increased RTs. Across clusters and frequency bands, good phase relations occurred in a correlated manner specifically when they processed the behaviorally relevant stimulus. Furthermore, the fronto-central cluster exerted a beta-band influence onto the occipital cluster whose strength predicted short RTs. These results suggest that local gamma and beta synchronization and their inter-regional coordination jointly improve behavioral performance.

## INTRODUCTION

Cognitive functioning depends on flexible communication among brain areas ^1–3^. This communication is likely subserved by rhythmic synchronization among distributed neuronal groups ^4, 5^, as proposed by the Communication-through-Coherence (CTC) hypothesis ^6, 7^. The CTC hypothesis states that brain rhythms entail phases of enhanced excitation and inhibition, respectively, and that inputs are particularly effective when they are consistently timed to avoid inhibition and align with excitation. This can be achieved if inputs are themselves rhythmic and entrain coherent rhythms in their target areas. Indeed, in the primate brain, rhythms in distinct frequency bands have been found to form distinct networks and to serve distinct roles ^8–19^. In particular, the gamma rhythm in the visual system shows entrainment, measured as Granger causality (GC), that is stronger in the bottom-up than top-down direction, while the beta rhythm shows the opposite pattern ^9, 10^. Moreover, these frequency-specific influences are interrelated such that top-down beta enhances bottom-up gamma ^20^. Thus, these rhythms are closely linked to the anatomically defined hierarchical order of visual areas, which reaches from early visual areas in occipital cortex, through mid-level areas in temporal and parietal cortex, to high-level areas in frontal cortex ^8, 10, 21, 22^.

Several studies have demonstrated the functional relevance of these rhythms, by linking them to behavioral performance ^12, 23, 24^. The reaction time (RT) of macaques to a stimulus change can be partly predicted by the strength of local gamma-band synchronization in area V4 at the time of the change ^25^. In fact, even the phase of the V4 gamma rhythm is similarly predictive of RT ^26^. The RT reflects the speed and/or efficiency with which the stimulus change is signaled from lower to higher visual areas and ultimately to motor-control areas, and thus the RT assesses inter-areal communication. This communication, according to CTC, should depend on the entrainment between relevant brain areas. Indeed, the visually induced gamma rhythm in macaque area V1 entrains area V4 at the phase relation that leads to shortest RTs; any momentary deviation from this typical gamma phase relation leads to longer RTs ^27^. In the present study, we investigated whether this holds for the gamma-band entrainment between occipital visual areas generally, whether it also holds for the beta-band entrainment between higher visual and fronto-central areas, and how those beta- and gamma-systems interact.

We found that indeed, gamma entrainment between all pairs of areas V1, V2, V4, and DP occurs at the phase relation preceding particularly short RTs. Intriguingly, also the beta entrainment among frontal and central areas S1, 5, F1, F2, and F4 occurs at the beta-phase relation preceding particularly short RTs. Finally, the beta synchronization between fronto-central areas correlated with the gamma synchronization between occipital areas. In fact, there is a directional influence, such that high fronto-central beta power precedes high occipital beta power. Furthermore, GC in the beta band is stronger in the fronto/central-to-occipital direction than vice versa, and the momentary precision of this beta GC is again partly predictive of RTs. These results support the notion that flexible communication depends on coordinated interareal synchronization of brain rhythms.

## RESULTS

Two macaques performed a selective visual attention task (see Figure 1A and Method details). Behavioral accuracy was far above chance level for both animals (Figure 1B). Behavioral RTs showed no difference between trials with attention to the right versus the left hemifield (Figure 1C; p = 0.16 for Monkey K, P=0.06 for monkey P).

**Figure 1.**
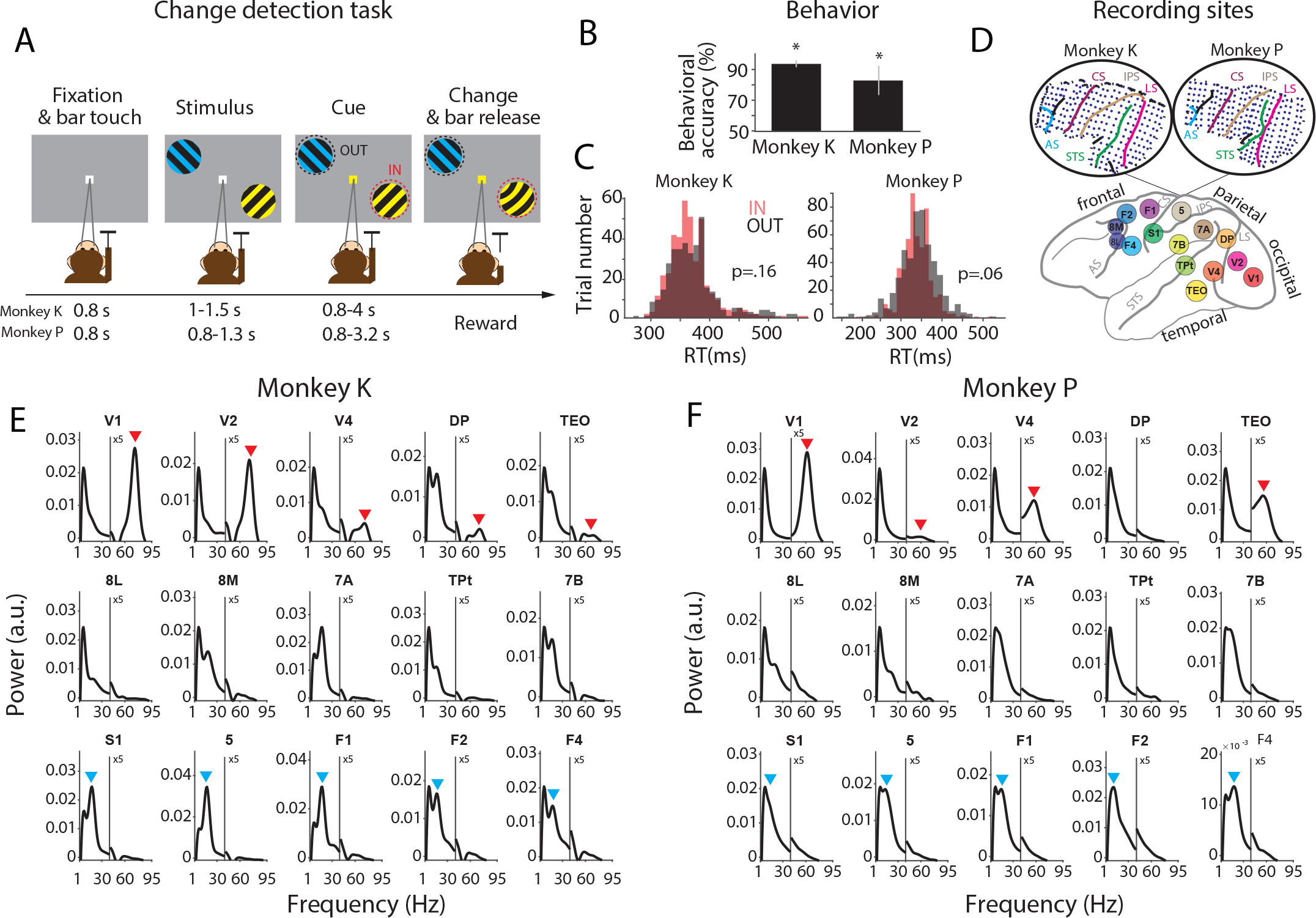
Stimuli and Task, performance, ECoG and power spectra. (A) Following a fixation period, two patches of grating with orthogonal orientations were presented. One stimulus was in the lower right visual quadrant, the other one was in the upper left quadrant at an equal eccentricity. One stimulus was tinted yellow, the other blue, with the colors randomly assigned across trials. Subsequently, the fixation point assumed the color of one of the stimuli, thereby cueing this stimulus to be the behaviorally relevant target stimulus, and leaving the other one to be the behaviorally irrelevant distractor stimulus. After a random delay of up to a few seconds, randomly either the target or the distractor underwent a small change. If changes in the target were reported by a bar release, a reward was given. If changes in the distractor were reported, a timeout was given. If changes in the distractor were not reported, they were always later followed by changes in the target, and if those were reported, a reward was given. (B) Behavioral accuracy (percentage of correct trials) per monkey. Error bars show SEM across sessions (9 and 14 sessions from Monkey K and P, respectively). Stars show significance with regard to the 50% chance level (z-test, *p* < 0.05). (C) Distributions of reaction times (RTs), separately for attend-IN (IN) and attend-OUT (OUT) conditions, per monkey. P-values are from a two-sample t-test between attention conditions (statistical t-test, p=0.16 and p=0.06 for monkeys K and P, respectively). (D) The ECoG covered 15 brain areas in occipital, temporal, parietal and frontal cortex with a total of 252 electrodes. Insets on top illustrate the location of electrodes in the array implanted in the left hemisphere of two monkeys. LS: lunate sulcus, STS: superior temporal sulcus, IPS: intraparietal sulcus, CS: central sulcus, AS: arcuate sulcus. (E, F) The average power spectra (after 1/f correction as explained in Methods) per area as indicated on top of each panel, averaged over all sessions, sites, and trials of the respective area. Spectra are shown separately for monkey K (E) and monkey P (F). The power spectra in the range of 1-95 Hz were calculated for the epoch from 200 ms to 0 ms before the stimulus change. Red and blue arrows indicate the gamma and beta peaks in occipital and fronto- central areas, respectively. Y-axes for the gamma band were multiplied by five, as indicated.

During multiple sessions with task performance, we used a chronically implanted electrocorticographic (ECoG) array to record from large parts of the left hemisphere. As the left-hemisphere visual areas have a selectivity for the contralateral right visual hemifield, trials with attention to the right hemifield are referred to as attend-IN condition, and trials with attention to the left hemifield as attend-OUT condition, abbreviated as IN and OUT conditions, respectively. A core behavioral benefit of attention is to shorten RTs. Therefore, we investigated whether RTs in response to the stimulus change could be predicted by rhythmic synchronization just prior to the change, in the epoch from 200 ms to 0 ms before the change (Rohenkohl *et al.* ^27^; see Method details).

The ECoG covered 15 brain areas in occipital, temporal, parietal and frontal cortex with a total of 252 electrodes (Figure 1D). From all electrodes simultaneously, local field potentials (LFPs) were recorded against a common reference. To remove the common reference, LFPs from immediately neighboring electrodes were subtracted from each other, and the resulting 218 bipolar derivations are referred to as “recording sites” (or just “sites”). When the sites were sorted according to their underlying brain area, the average power spectra per area (after removing the 1/f component) showed distinct peaks. In particular, occipital areas V1, V2, V4, DP and TEO showed a gamma peak, and fronto-central areas S1, 5, F1, F2 and F4 showed a beta peak (Figures 1E and 1F). For each pair of areas, there were multiple site pairs (range: 4 – 1296). For each individual site pair, rhythmic synchronization was quantified using the pairwise-phase-consistency (PPC) metric ^28^.

### Attention induces interareal synchronization at optimal phase relation

To investigate a putative link between synchronization and behavioral RT, we used an approach developed by Rohenkohl *et al.* ^27^ for V1-V4, modified it, and expanded it to more areas. We first illustrate our modified approach for two example area pairs, V1-V4 and F1-F4.

The complete set of all PPC spectra for these example area pairs is shown in Figure S1. This reveals a substantial variability of synchronization across site pairs, likely due to stronger synchronization between site pairs with stronger connectivity ^8^. We selected, for further analysis, the site pairs whose PPC exceeded, at any frequency, a threshold. The threshold was obtained by taking all PPC values over all site pairs of all area pairs, determining their mean and SD, and defining the threshold as the mean+3SD.

The interareal PPC spectra averaged over the selected site pairs were dominated by a gamma peak for the area pair V1-V4 (Figures 2A and 2C) and by a beta peak for the area pair F1-F4 (Figures 2B and 2D). These peaks were present in both animals, with slightly different individual peak frequencies. We determined the individual gamma and beta peak frequencies as the frequency with the highest average PPC in the respective frequency band, and we aligned the further analyses to those individual peak frequencies (±15 Hz for beta, ±20 Hz for gamma).

**Figure 2.**
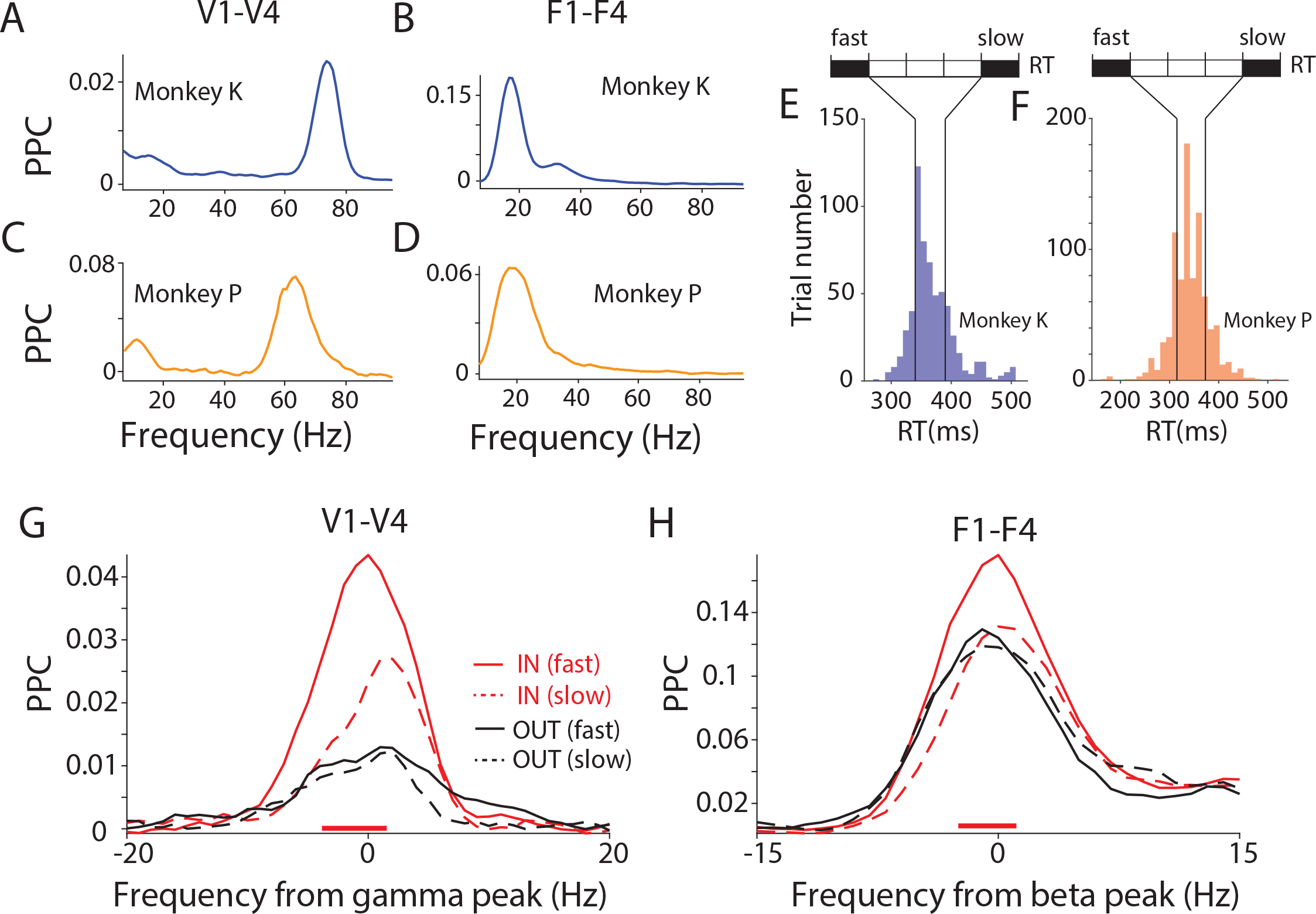
Fast reaction times are associated with higher PPC at dominant frequency. (A, B) PPC averaged over selected site pairs of V1-V4 as a sample occipital area pair (A), and F1-F4 as a sample fronto-central area pair (B), for monkey K. (C, D) The same as (A, B) but for monkey P. (E, F) Distributions of RTs for monkey K (E) and monkey P (F). Based on RTs, trials were sorted and then separated into five bins with equal number of trials. The first and last bins were defined as fast-RT and slow-RT trials, respectively, and used for the respective comparisons. (G) Average V1-V4 PPC modulation in fast-RT trials (solid lines) and slow-RT trials (dashed lines), separately for conditions IN (red) and OUT (black), averaged over both monkeys. The PPC close to the gamma peak was higher for fast- than slow-RT trials, specifically during the IN conditions, as indicated by the red horizontal line on the bottom. (H) The same as (G) but for the F1-F4 PPC, aligned to beta peak. See also Figure S1.

As a first test for a relation between PPC and RT, we compared PPC between trials with fast versus slow RTs. The RTs of each monkey separately were sorted and divided into five equal bins (Figures 2E and 2F). The first, fast-RT, and the last, slow-RT, bin were compared. For the trials in those two bins, the PPC was first averaged over all site pairs per monkey and then over the two monkeys. PPC between V1 and V4 was stronger before fast versus slow RTs at the gamma peak (Figure 2G; non-parametric randomization test with correction for multiple comparisons, see Methods for details), confirming a related previous analysis ^27^. Importantly, a very similar effect existed also for PPC between F1 and F4 at the beta peak (Figure 2H). Both for the V1-V4 gamma and the F1-F4 beta effect, this effect was only present in the IN condition, when the contralateral visual stimulus was behaviorally relevant. The effect was absent in the OUT condition, when the behavioral response was to the ipsilateral stimulus, and the contralateral stimulus was behaviorally irrelevant. This demonstrates that the effect is spatially specific and thereby most likely not simply reflecting RT fluctuations due to arousal fluctuations.

Next, we expanded our investigation of the relationship between neuronal synchronization and behavioral RT to further area pairs. For the areas of the visual system (V1, V2, V4, TEO, DP, 7a, 8L, 8M) coherence between all area pairs shows a beta peak, and coherence between most area pairs also shows a gamma peak, as we have shown previously for this dataset ^10^. In the present study, we focused on those area pairs showing the strongest respective peaks. As illustrated for V1-V4 and F1-F4, PPC spectra were averaged over selected site pairs, separately for all area pairs, including the non-visual areas Tpt, 7B, S1, area 5, F1, F2, F4 (Figure S2). The average coherence varied substantially across area pairs, both for gamma (Figure S2A) and for beta (Figure S2B), likely due to stronger synchronization between area pairs with stronger connectivity ^8^. We selected, for further analysis, the area pairs whose average PPC exceeded, at any frequency, a threshold. The threshold was obtained by taking all average PPC values over all area pairs, determining their mean and SD, and defining the threshold as the mean+2SD. This resulted in one cluster of neighboring areas per frequency band: Gamma PPC was particularly strong in a cluster of occipital areas consisting of areas V1, V2, V4, and DP (Figure S2A, dashed box); Beta PPC was particularly strong in a cluster of fronto-central areas consisting of areas S1, 5, F1, F2, and F4 (Figure S2B, dashed box). The further analyses focus on the synchronization within and between those clusters and its relation to selective attention and behavioral RT.

For the occipital cluster (V1, V2, V4, and DP, Figure 3A), the average inter-areal PPC spectrum shows a gamma PPC peak that is stronger during the IN than the OUT condition (Figure 3B, Wilcoxon ranksum test, *p* < 0.05; Bonferroni-corrected for multiple comparisons across frequencies). The distribution of gamma-peak PPC values across all selected inter-areal site pairs of the occipital cluster showed significantly stronger gamma for the IN than the OUT condition (Figure 3C; t-test, *p* < 0.05). Attention shortens behavioral RTs, and we therefore tested whether single-trial RTs can be (partly) predicted by the precision of single-trial inter-areal gamma phase locking. We followed the approach developed by Rohenkohl *et al.* ^27^, which we illustrate first for one gamma-synchronized V1-V4 site pair (Figure 3D-H): (1) Per site pair and per trial, the gamma phase relation is estimated (Figure 3D); (2) The average phase relation over trials is determined, and all single-trial phase relations are rotated such that the average phase relation is zero (Figure 3F); (3) After this rotation, the single-trial phase relations reflect deviations from the phase relation at which the two sites synchronize on average; we define the cosine of these deviations to be the Goodness of Phase Relation (GPR); The GPR assumes a value of one for single-trial phase relations equal to the average, and a value of minus one for single-trial phase relations opposite to the average; (4) The correlation between GPR and RT is calculated (Figure 3H).

**Figure 3.**
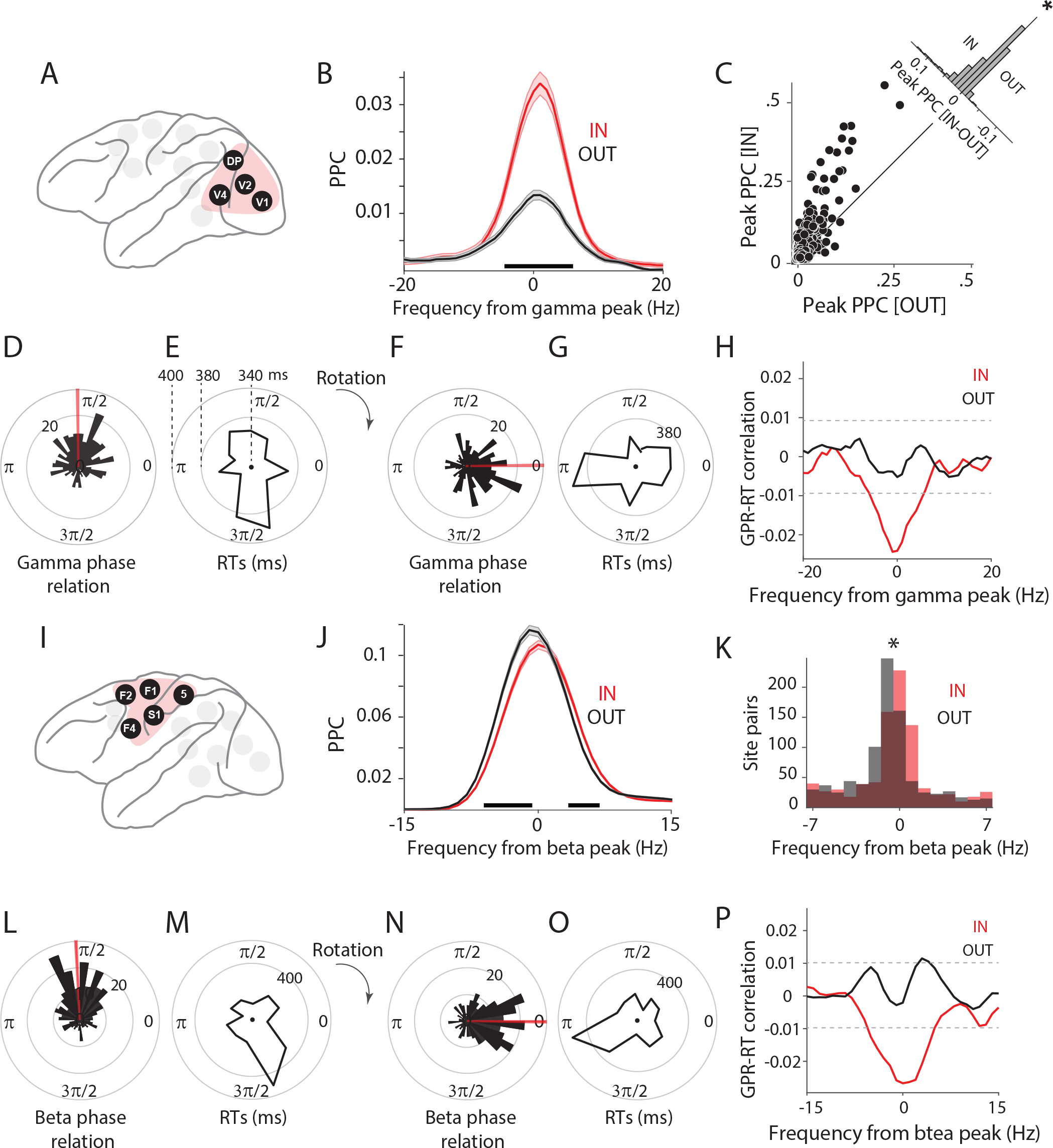
Phase relation at dominant frequency predicts behavior. (A) The occipital cluster: V1, V2, V4, DP. These areas showed particularly strong PPC in the gamma band. (B) Average PPC between area pairs in the occipital cluster, for IN (red) versus OUT (black) conditions. Shaded areas indicate SEM across site pairs. Black horizontal line indicates frequencies with a significant difference between IN and OUT conditions. (C) Scatter plot of peak PPC strength for IN vs. OUT condition. Each dot represents an interareal site pair from the occipital cluster. The inset shows the histogram of differences (IN - OUT) between peak PPC strength. (D) Distribution of gamma phase relations between an example V1-V4 site pair, across trials. Red line shows the mean phase of this site pair. (E) Distribution of RTs as a function of V1-V4 gamma phase relation. (F and G) All phase relations (F) and their corresponding RTs (G) were rotated to bring the mean phase to zero. (H) Correlation between goodness of phase relations (GPRs) and RTs, as a function of frequency for IN (red) and OUT (black) conditions. Black horizontal dashed lines indicate significance thresholds, corrected for multiple comparisons. (I) The fronto-central cluster: S1, 5, F1, F2, F4. These areas showed particularly strong PPC in the beta band. (J) Same as (B), but for fronto-central areas and aligned to the beta peak. (K) Histogram of peak PPC frequencies relative to the beta peak frequency. The star denotes a significant difference between IN and OUT distributions (t-test, *p*<0.05). (L-P) Same as (D-H), but for fronto-central areas and aligned to the beta peak. See also Figures S2 and S3.

Single-trial phase relations showed a uni-modal distribution, reflecting synchronization at a preferential phase relation (Figure 3D). Importantly, this preferential phase relation showed relatively short RTs, whereas the opposite phase relation showed longer RTs (Figure 3E). The rotation of the average phase relation to zero directly illustrated increased RTs for deviations from the average phase relation (Figures 3F and 3G). The GPR-RT correlation revealed a significant gamma peak, specifically for the IN condition (Figure 3H; non-parametric randomization test, p < 0.05, corrected for multiple comparisons). The peak was negative, indicating that the average phase relation was related to short RTs. This peak was absent in the OUT condition (Figure 3H). Thus, the effect described previously by Rohenkohl *et al.* ^27^ for V1-V4 holds when all areas of the occipital cluster with particularly strong gamma PPC are combined.

Next, we investigated whether a similar effect was also present for the beta PPC in the fronto-central cluster (S1, area 5, F1, F2, and F4, Figure 3I). For the example frontal area pair F1- F4, we have already demonstrated above that the beta PPC is stronger for trials with fast RT compared to slow RT (Figure 2H). Therefore, we repeated the analysis of attention effects and of single-trial phase relations and RTs for the entire cluster (Figures 3I-3P). The inter-areal PPC spectrum averaged over all selected inter-areal site pairs in this cluster of areas revealed that attention primarily shifted the beta peak to a slightly higher frequency. This leads to PPC decreases on the rising, and PPC increases on the falling flank of the beta peak (Figure 3J; Wilcoxon ranksum test, *p* < 0.05; Bonferroni-corrected for multiple comparisons across frequencies). The distribution of beta-peak frequency values across all selected inter-areal site pairs of the fronto-central cluster showed significantly higher beta-peak frequencies for the IN than the OUT condition (Figure 3K; t-test, *p* < 0.05). The example F1-F4 site pair showed a uni-modal distribution of single-trial beta phase relations (Figure 3L). The average phase relation was associated with relatively short RTs, whereas the opposite phase relation was associated with long RTs (Figures 3L-3O). The GPR-RT correlation showed a significant negative beta peak for the IN condition (Figure 3P; non-parametric randomization test, p < 0.05, corrected for multiple comparisons). For the OUT condition, two adjacent frequency bins slightly above the alignment frequency reached significantly positive values.

These correlations of RT with GPR are likely not due to a correlation of RT with LFP power. The power-RT correlation spectra for occipital areas in the gamma band showed no significant results (Figure S3A), and for fronto-central areas in the beta band showed some marginally significant frequency bins that did not match the GPR-RT correlation spectrum (Figures S3B, 3P).

### GPR-RT correlation increases by averaging over simultaneous recording sites

The GPR-RT correlation values that we report here are based on GPR and RT measurements for single trials, which incurs relatively much measurement and/or estimation noise. Such noise leads to an underestimation of the true correlation. However, the true correlation cannot be recovered by eliminating noise via binning of trials and averaging GPR and RT within those bins before calculating the correlation; rather, this procedure arbitrarily inflates the estimated correlation value ^20, 29^. To avoid this inflation and still eliminate noise and come closer to the true trial-by-trial correlation, we averaged single-trial GPR values over site pairs before calculating the GPR-RT correlation. We first averaged GPR values over all site pairs per area pair, and then used these area-pair GPRs to determine their correlation with RT. This GPR- RT correlation, during the IN condition, showed a significant negative gamma peak for the occipital areas (Figure 4A), and a significant negative beta peak for the fronto-central areas (Figure 4B). Intriguingly, the correlation values for area-pair GPRs were approximately fourfold larger than for site-pair GPRs (Figure 4C). Therefore, we further averaged GPR values over all site pairs per cluster of areas, and then used these cluster GPRs to determine their correlation with RT. The resulting correlation values were approximately sixfold larger than for site-pair GPRs (Figure 4C). Finally, we averaged GPR values over all site pairs per cluster of areas, and additionally over the two clusters; note that this entails averaging of GPR over two frequency bands, that is, gamma for the occipital and beta for the fronto-central cluster. This resulted in a GPR-RT correlation that was approximately eightfold larger than for single site pairs. Together, these results suggest (1) that the true single-trial correlation is substantially stronger than estimated on the basis of single site pairs, (2) that GPRs within an area pair and within a cluster are correlated such that noise can be eliminated by averaging over them, and (3) that this correlation between GPRs also holds between occipital gamma GPRs and fronto-central beta GPRs.

**Figure 4.**
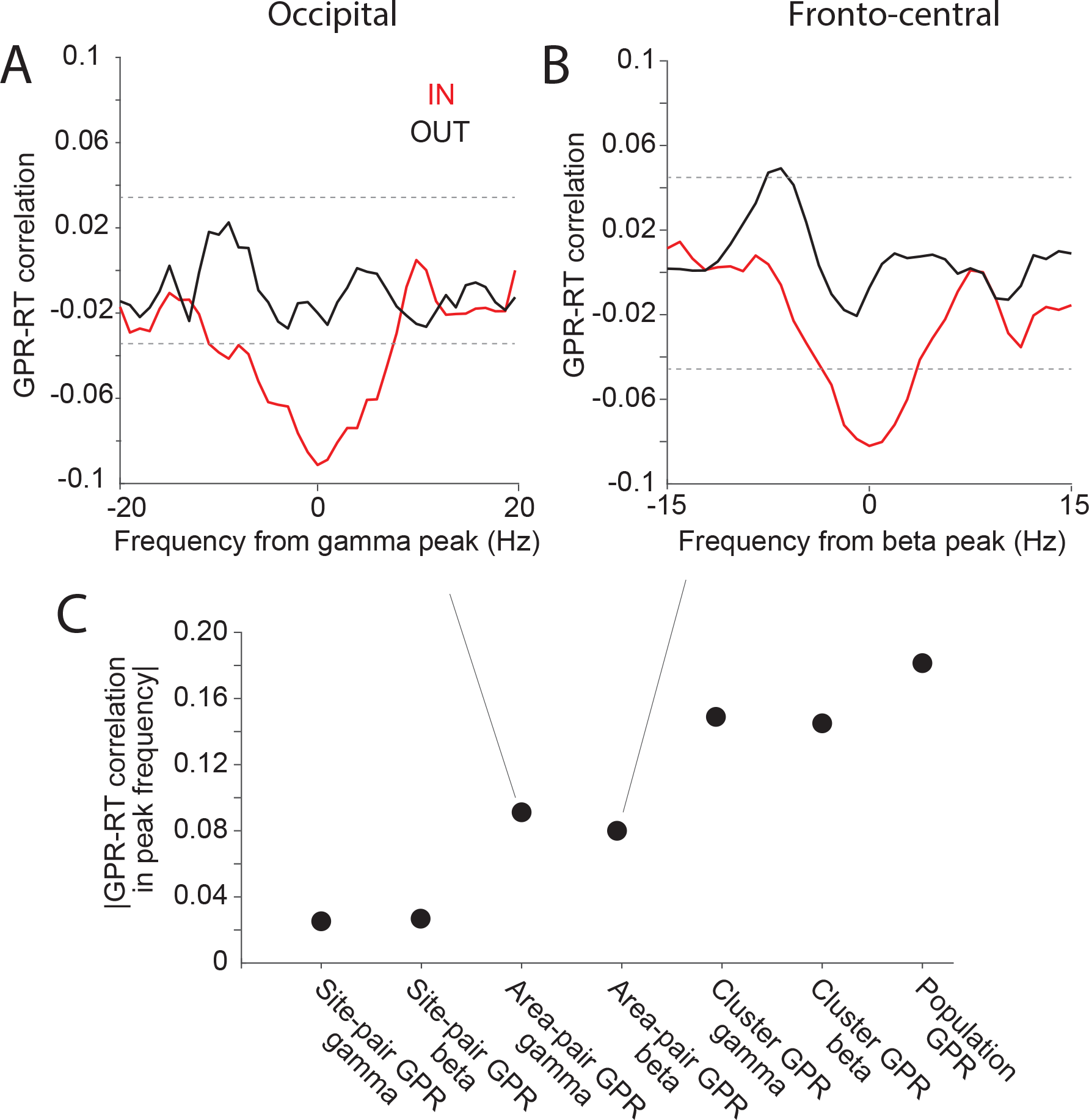
GPR averaging over a population of recording sites reduces the noise effect and enhances the correlation values. (A-B) Correlation between GPRs and RTs across trials, after first averaging GPRs over site pairs of an area pair (defined as area GPR), before calculating the GPR-RT correlation. This GPR-RT correlation is shown for the occipital cluster aligned to the gamma peak (A), and for the fronto-central cluster aligned to the beta peak (B). (C) GPR-RT correlation based on GPRs at the level of site pairs (corresponding to Figure 3H, 3P), area pairs (Figure 4A, 4B), clusters (average GPRs across all site pairs in each cluster), and the complete population (average GPRs across all site pairs of two clusters combined over the respective dominant frequency bands).

### Occipital gamma GPRs correlate with fronto-central beta GPRs

We investigated this latter point directly, by testing for a correlation between occipital cluster-level gamma GPRs and fronto-central cluster-level beta GPRs. This correlation was present during the IN conditions (Figure 5A, r = 0.19, p=7.3x10^−8^, Pearson correlation), but not during the OUT conditions (Figure 5B, r = 0.06, p=.067). The observed difference between the IN and the OUT conditions was significant (Figure 5C, p<0.05, z-test comparing observed difference to a randomization distribution obtained after randomly permuting trials across conditions). Thus, fronto-central beta and occipital gamma GPRs fluctuate across trials in a coordinated manner, suggesting some link between these regions in their dominant rhythms.

**Figure 5.**
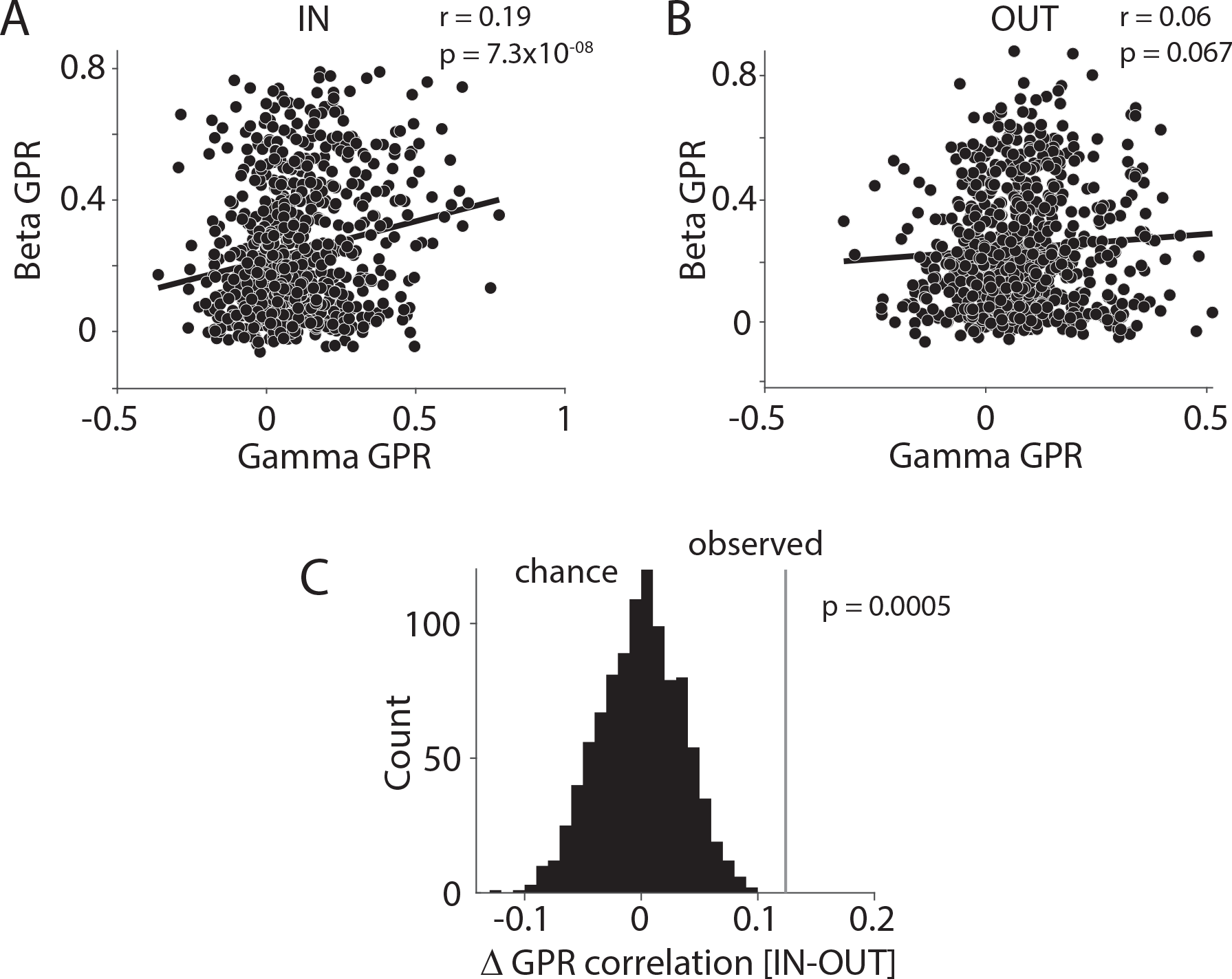
Occipital gamma GPR correlates with fronto-central beta GPR during IN condition. (A, B) Across-trial correlation between occipital gamma GPR and fronto-central beta GPR, separately for condition IN (A) and OUT (B). Each dot represents the respective GPR values from one trial, averaged over site pairs of the corresponding areas (gamma GPR in occipital areas and beta GPR in fronto-central areas). (C) Comparison of the empirically observed difference in GPR correlation (IN-OUT) with chance distribution based on 1000 randomizations of trials. See also Figures S4.

### Directed inter-areal influences as assessed by Granger causality predict RT

Next, we complemented the analysis of GPR with an analysis of a metric of directed inter-areal influences, namely Granger causality (GC). The GC analysis used the selection of site pairs and area pairs based on PPC; a selection based directly on GC gave almost the same selected site pairs, or area pairs, respectively. GC between occipital areas in the gamma band was stronger in the bottom-up than the top-down direction, and was increased by attention, in line with previous reports (Figure 6A, Wilcoxon ranksum test, *p* < 0.05; Bonferroni-corrected for multiple comparisons across frequencies) ^10, 20, 30, 31^. Interestingly, the correlation between occipital gamma GC and RT showed a clear negative peak specifically during the IN condition and in the bottom-up direction (Figure 6B, non-parametric randomization test, p < 0.05, corrected for multiple comparisons). This correlation was calculated across single trials by using the jackknife correlation approach ^29^. GC between fronto-central areas in the beta band was stronger in the bottom-up than top-down direction (Figure 6C, Wilcoxon ranksum test, *p* < 0.05; Bonferroni-corrected for multiple comparisons across frequencies). Note that beta GC between areas of the visual hierarchy has been shown to be stronger in the top-down direction ^9, 10^. Potential reasons for this difference between the visual system and fronto-central regions will be explored in the discussion. The correlation between fronto-central beta GC and RT showed negative peaks specifically during the IN condition and most pronounced in the dominant, bottom-up, direction (Figure 6D, non-parametric randomization test, p < 0.05, corrected for multiple comparisons).

**Figure 6.**
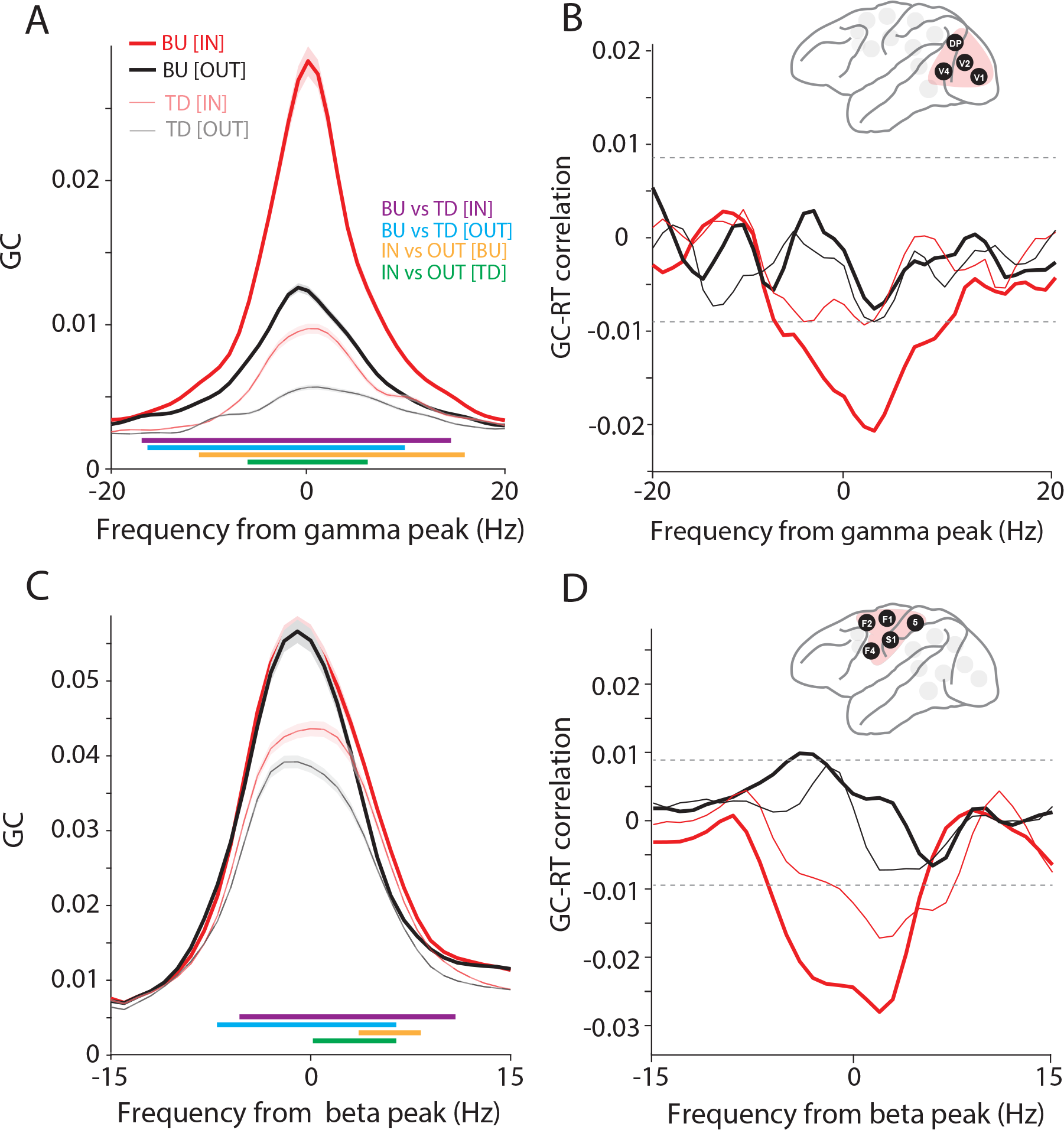
Occipital gamma and fronto-central beta Granger causality predict RTs. (A) Interareal Granger causality (GC), averaged over all site pairs of the occipital cluster, aligned to the gamma peak, separately in the bottom-up (BU, tick lines) and top-down (TD, narrow lines) directions, and for the IN (red) and OUT (black) conditions. Colored horizontal lines denote significant differences between conditions (IN, OUT, BU, TD), as indicated in the color legend. (B) Jackknife correlation (see Methods for details) between single-trial GCs and RTs. The inset shows the selected areas in the occipital cluster. (C, D) Same as (A, B), but for areas in the fronto-central cluster, and aligned to the beta peak.

Fronto-central beta GC values and occipital gamma GC values showed a positive correlation across single trials during the IN condition (r = 0.11, P=0.0026, Pearson correlation), but not during the OUT condition (r = -0.05, p=0.26), and this attentional effect was significant (Figure S4). This is similar to the above mentioned related analysis for GPR.

### Directed influences from fronto-central to occipital areas and their behavioral relevance

Finally, we investigated the influences between the two clusters of fronto-central and occipital areas, and we tested whether any influences have behavioral relevance. Such long-range influences can be assessed with high sensitivity by means of power-power correlation, sometimes also referred to as amplitude envelope correlation ^32^. The time-varying power of beta and gamma were estimated for the last 400 ms before the behaviorally relevant stimulus change, using 200 ms windows shifted in 50 steps of 4 ms (Figure 7A). The Pearson correlation coefficient was calculated between power time courses, as a function of lag, pooling data points from all trials. This was done for all four combinations of clusters (fronto-central or occipital) and rhythms (beta or gamma). Only one combination showed a significant correlation, namely fronto-central beta versus occipital beta (Figure 7B). This correlation showed a lagged peak indicating that fronto-central beta was leading occipital beta by 32 ms.

**Figure 7.**
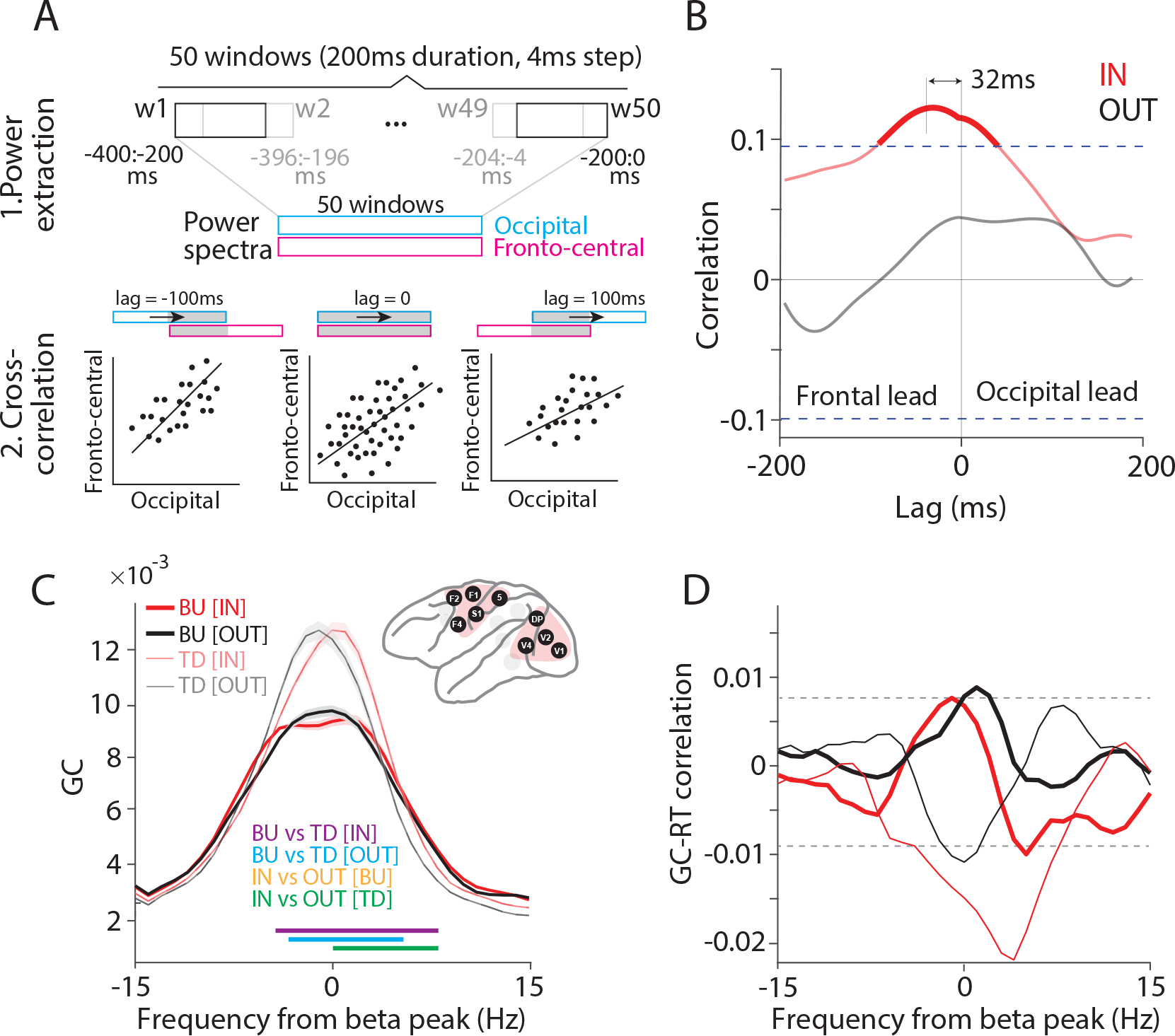
Fronto-central and occipital areas communicate through top-down beta influences. (A) Schematic illustration of time-lagged cross-correlation between fronto-central and occipital areas. The power spectra were calculated, separately for the occipital and the fronto-central cluster, for 50 windows with a duration of 200 ms and a shift of 4 ms. The first window is from -400 ms to -200 ms, the 50^th^ window from -200 ms to 0 ms relative to the stimulus change. The correlation between the two sets of power spectra is calculated in two steps: (1) Calculating power spectra. (2) Calculating cross-correlation across trials, as a function of lag between windows, across the windows that overlap for a given lag. (B) Time-lagged cross-correlation between beta power (14-16 Hz) of fronto-central and occipital areas, separately for IN (red) and OUT (black) conditions. Dashed lines show significance thresholds based on the randomization approach and corrected for multiple comparisons. (C) GC between fronto-central and occipital clusters aligned to beta peak, for IN (red) and OUT (black) conditions, and in bottom-up (BU, tick lines) and top-down (TD, narrow lines) directions, separately. Colored horizontal lines indicate significant frequency bands for the indicated comparisons. (D) Jackknife correlation (see Methods for details) between single-trial RTs and beta GCs between the occipital and fronto-central cluster for IN (red) and OUT (black) conditions and in the bottom-up (BU, tick lines) and top-down (TD, narrow lines) directions.

Therefore, we further investigated the interactions between the fronto-central and the occipital cluster at beta. As for the inter-areal analyses within the clusters, we now selected all inter-cluster site pairs whose PPC exceeded the mean+3SD over all respective PPC values. We first tested whether the beta GPR was related to RT, and found no relation. We then investigated the GC between the two area clusters and found that it was stronger in the top-down than bottom-up direction, i.e., stronger from the fronto-central to the occipital cluster than vice versa (Figure 7C). Interestingly, during the IN condition, the beta peak shifted to a slightly higher frequency (Figure 7C; green horizontal lines indicating significant increases in falling flank). The trial-by-trial correlation between beta GC and RTs showed a prominent negative peak for the top-down direction during the IN condition (Figure 7D; non-parametric randomization test, p < 0.05, corrected for multiple comparisons). This suggests that top-down beta influences from fronto-central areas to occipital areas increase the speed and/or efficiency with which the behaviorally relevant stimulus change is communicated between brain areas and ultimately turned into a behavioral response.

### Purely frontal cluster shows similar effects as fronto-central cluster

We investigated whether the core findings for the fronto-central cluster also held when we restricted it to only contain frontal areas, that is, areas anterior to the central sulcus. The following figures, obtained for the frontal cluster, correspond to the figures listed in parenthesis, obtained with the fronto-central cluster: Figure S5A,D (Figure 3J,P), Figure S5B,E (Figure 6C,D), Figure S5C,F (Figure 7C,D). While there are differences in details, the results are qualitatively similar.

## DISCUSSION

In summary, we found interareal synchronization to be particularly prominent among occipital areas in gamma, and among fronto-central areas in beta. In both clusters of areas, and for both frequency bands, interareal synchronization occurred at the phase relation that led to the shortest reaction times, and deviations from that phase relation led to systematically slower reaction times. Both for occipital gamma and fronto-central beta, similar results were obtained for interareal GC, with stronger GC leading to shorter reaction times. Also, stronger GC from the fronto-central to the occipital cluster at beta led to shorter reaction times. Occipital gamma GPR and fronto-central beta GPR showed trial-by-trial correlation, specifically during the IN condition; the same held for GC. Together, these findings suggest that all these interareal synchronization and entrainment phenomena improve behavioral performance. The effects were mostly specific to the IN condition, indicating that they did not reflect unspecific arousal changes. Also, the effects could not be explained by corresponding changes in power. The effects found for the fronto-central cluster remained similar when the cluster was restricted to the frontal areas.

The trial-by-trial prediction of RT by GPR improved fourfold when GPR was averaged over site pairs within area pairs, sixfold when averaging over area pairs within clusters, and eightfold when averaging over clusters and frequency bands. This indicates (1) that individual site pairs provide only noisy estimates of interareal synchronization and entrainment, (2) that GPR is correlated across site pairs, area pairs and even the two frequency bands, and (3) that the noisy GPR estimates lead to an underestimation of its true predictive power for behavioral reaction times.

This study is based on recordings from two macaques, as is typical for awake macaque neurophysiology. With two animals, any useful inference is limited to the investigated sample ^33^. Furthermore, this study is based on LFP data. The LFP, in the frequency ranges investigated here, primarily reflects postsynaptic potentials and thereby neuronal inputs. Yet, ≈80% of synaptic inputs to a cortical neuron are generated from locally neighboring neurons through their spiking ^34^. Thus, local neuronal spiking is probably the main source of the LFP ^35–37^. Nevertheless, it has been argued that the LFP also reflects synaptic inputs from other areas, and that this explains interareal coherence and GC ^38^. This mechanism predicts that interareal coherence and GC directly reflect power in the sending area. Correspondingly, power should be equally or more predictive of behavior than interareal coherence and GC. We find the opposite: While both interareal coherence and GC are predictive of behavior, power is not. Similar results have been reported before, e.g., cortico-muscular coherence reflects the hazard rate and thereby RTs, and this holds even when the data are stratified for power ^39^. At the same time, LFP provides major advantages for investigating interareal rhythmic synchronization. Rhythmic synchronization between two areas can only be properly quantified if the local rhythmic activities are measured both sensitively and independently. The sensitivity of the LFP in this regard is better than that of spike recordings, because spike probability is modulated by the rhythm’s phase only to some degree, and spikes sample the rhythmically modulated spiking probability only at very few discrete time points. The independence of LFP recordings from two areas is better than that of EEG or MEG recordings, because they suffer from signal mixing even after source projection ^40–42^, although much progress has been made to address this ^43–45^. Thus, between the macroscopic EEG/MEG and the microscopic spike recordings, the mesoscopic LFP occupies a sweet spot for assessing interareal coherence and GC. This assessment still requires the removal of the common recording reference, e.g., through bipolar derivation as done here, or through using Laplacian operators ^46^.

We used simultaneous bipolar LFP recordings from many areas to investigate whether interareal coherence and GC play functional roles for interareal communication. We evaluated the efficiency of this communication by measuring the behavioral reaction time, i.e., the time that the go signal took to travel from the retina through the different cortical areas to motor cortex and spinal cord to finally issue the behavioral response. Behavioral RTs can be partly predicted by the local neuronal gamma synchronization in macaque area V4 ^25, 27^. Also, the gamma power in the human middle occipital gyrus predicts RTs when investigated with source-projected MEG ^47^. Such MEG source power estimates reflect neuronal synchronization both within and across neighboring areas. Indeed, the coherence between macaque areas V1 and V4 is enhanced before short RTs ^27^. Crucially, this study also showed that V1-V4 coherence occurs at the phase relation leading to the shortest RTs and thereby improving behavioral performance. Here, we built on this approach and expanded it to all areas covered by the ECoG, leading to the described findings in fronto-central regions and in the beta band.

Rhythmic activity in fronto-central regions has previously been related to motor performance. Reaction times in a simple visuomotor reaction-time task are correlated with gamma-band activity before the go cue measured with source-projected EEG from human fronto-parietal areas ^48^. Also, human subjects show a correlation between their readiness to respond and their coherence between motor cortex and spinal cord, which is positive for the gamma band and negative for the beta band ^39^. Furthermore, several studies have demonstrated that strong motor-cortical beta activity was predictive of slower movements, i.e. movements with lower peak acceleration, yet beta was not related to reaction times ^49^. Similarly, when transcranial alternating current stimulation (tACS) was applied either in the beta band, at 20 Hz, or in the gamma band, at 70 Hz, movement speed was either decreased or increased, respectively ^50, 51^. In summary, these studies suggest that motor cortical gamma is involved in promoting new movements, whereas beta is involved in stabilizing the current motor state or posture ^52^.

We found that fronto-central beta synchronization and fronto/central-to-occipital beta GC is predictive of short RTs. This is in line with a previous analysis of the same dataset, showing that moment-to-moment enhancements of top-down beta GC from area 7a onto V1 lead to corresponding enhancements of bottom-up gamma GC from V1 onto V4 ^20^. It might be relevant that these and the present results were obtained with a change detection task. In this task, the current state of the stimulus needs to be constantly compared to the most recent state of the stimulus, which is likely provided by top-down signaling, which in turn is related to interareal GC in the beta band ^8–10^. More generally, our findings are in line with a review of the beta literature concluding that “beta oscillations observed in sensorimotor cortex may serve large-scale communication between sensorimotor and other areas and the periphery” ^53^.

We have previously analyzed GC between a subset of the areas studied here, namely between the areas of the visual system: V1, V2, V4, TEO, DP, 7A, 8L and 8M. For these areas, anatomical studies in macaques have shown that interareal laminar projection patterns largely abide by a global hierarchy that assigns a hierarchical level to each area ^21, 54^. We previously found that between those areas, GC in the gamma band is typically stronger in the bottom-up than top-down direction, whereas GC in the beta band is typically stronger in the top-down than bottom-up direction ^10^. A highly similar pattern was found in a cohort of 43 human subjects studied with source-projected MEG ^9^. In the present study, we found that gamma GC among occipital areas is predictive for RTs, specifically for the gamma GC in the bottom-up direction (Figure 6B). Furthermore, we found that beta GC between fronto-central areas is predictive of RTs, for GC in both directions, yet stronger for the bottom-up direction; beta GC between strictly frontal areas, excluding post-central areas, is predictive of RTs, specifically in the bottom-up direction. These latter results, linking RT to beta GC predominantly in the bottom-up direction might appear surprising, given that beta-GC is stronger in the top-down direction between areas of the visual system. Yet, between the recorded frontal areas, i.e., F1, F2 and F4, beta GC is stronger in the anatomically defined bottom-up direction (Figure S6). Importantly, the bottom-up direction corresponds to the direction from the motor cortex to premotor areas. Thus, between these areas, the bottom-up direction arguably corresponds to the direction of functional feedback in the sense of corollary discharges ^55^. Therefore, we would like to speculate that beta is generally stronger in this direction of functional feedback, which in sensory systems is top-down, and in the (pre-)motor system is bottom-up. Note that this also holds for the other recorded sensory system, namely the somatosensory system.

Between the recorded somatosensory areas, i.e., S1 and area 5, beta-band GC was stronger in the top-down than the bottom-up direction (Figure S6). Between the different sensory systems, and/or between them and the (pre-)motor system, hierarchical relationships are hard to interpret, and we therefore refrain from that.

In conclusion, our analysis lends further support that behavioral performance is subserved by gamma synchronization between occipital areas, and importantly also by beta synchronization among frontal or fronto-central areas, and by the beta-band GC from frontal/fronto-central to occipital areas. These effects were not explained by the power of the respective rhythms, suggesting that they were genuine effects of interareal synchronization.

## EXPERIMENTAL MODEL AND SUBJECT DETAILS

All experimental procedures were approved by the ethics committee of Radboud University Nijmegen (Nijmegen, the Netherlands). Parts of the data have been used in other publications, e.g., ^8, 10, 20, 27, 56, 57^. The procedures and paradigms are described in ^10, 30^. Here we provide further details necessary for understanding the present analysis.

## METHOD DETAILS

### Behavioral Task and Electrophysiological Recording

Two adult male macaque monkeys (Macaca mulatta) were trained in a visual attention task. Throughout the task, the monkeys were required to maintain their gaze on a fixation point at the center of the screen. Each trial started when the monkey pressed the lever and fixated on the fixation point. Following an 800 ms fixation period, two isoluminant and isoeccentric stimuli appeared on the screen. Each stimulus was a drifting sinusoidal grating (diameter: 3 degrees visual angle; spatial frequency: ≈1 cycle/degree; drift velocity: ≈1 deg/s; temporal frequency: ≈1 cycle/s; contrast: 100%). Stimuli were controlled using the CORTEX software (http://www.cortex.salk.edu) and presented on a cathode ray tube (CRT) monitor at a refresh rate of 120 Hz. In each trial, yellow and blue tints were assigned randomly to the bright grating stripes of the two stimuli. The tints were present for the entire duration of stimuli presentation. After a variable stimulus period (1000-1500 ms in Monkey K, 800-1300 ms in Monkey P), the color of the fixation point changed to blue or yellow, cueing the stimulus with the corresponding tint to be the behaviorally relevant, attended, target, leaving the other one to be the behaviorally irrelevant, unattended, distractor. Transient shape changes (grating stripes undergoing a gentle bend, lasting 150 ms) of any one of the stimuli (target or distractor) could occur already before cue onset and until 4500 ms after cue onset, and occurred equally likely in the target and distractor. The monkey was rewarded for releasing a lever shortly (within 150-500 ms) after a change of the target, while ignoring changes of the distractor. All trials included a change of the target, either as the first change or after a distractor change. Trials during which the monkey broke fixation or released the bar outside the response window terminated without reward. Both monkeys performed the task with an accuracy far above chance (accuracy of 94% and 84% for monkeys K and P, respectively). Trials with attention directed to the stimulus in the visual hemifield contralateral (ipsilateral) to the recorded hemisphere are referred to as attend-IN (attend-OUT) condition, abbreviated as IN (OUT) condition.

During the task, neuronal activity was recorded from subdural ECoG grids realizing simultaneously 252 recording electrodes over the left brain hemisphere for a total of 15 brain areas (1 mm electrode diameter and 2-3 mm space between electrodes). The 15 areas were hierarchically ordered. Visual areas were ordered according to Bastos *et al.* ^10^ and Markov *et al.* ^21^; somatosensory and motor areas were ordered according to Vezoli *et al.* ^8^ and Chaudhuri *et al.* ^22^.

Signals were amplified by eight 32-channel Plexon headstage amplifiers (Plexon, USA), referenced against a silver wire implanted epidurally over the right occipital cortex (common recording reference). Signals were then filtered between 0.159 Hz and 8 kHz and digitized at approximately 32 kHz with a Digital-Lynx acquisition system (Neuralynx, USA). LFPs were obtained by low-pass filtering at 250 Hz and down sampling to 1 kHz. During the experiment, monkeys’ eye position was monitored by a video-based eye-tracking system with 230 Hz sampling rate (Thomas Recording, Giessen, Germany).

### Data Analysis

All analyses were carried out in MATLAB 2020b (The MathWorks, Inc., Natick, Massachusetts, USA) using the FieldTrip toolbox (http://www.fieldtriptoolbox.org/) ^58^ and custom MATLAB scripts.

Signals were re-referenced via a bipolar scheme by differentiating neighboring electrodes on the same lane of the ECoG grid, which resulted in 218 recording sites. This process improves signal localization, cancels the common recording reference, and rejects noise specific to each headstage. We use the term “electrode” to refer to a single unipolar recording electrode, and the term “site” to refer to a local bipolar derivation.

Using a discrete Fourier transform, 50 Hz line noise and its harmonics (100 and 150 Hz) were removed from signals. After removing trials with artifacts (trials with a variance five times greater than the mean variance in the same site), each trial was linearly detrended, which entails a demeaning. Within each recording site, the signal was normalized (subtract mean and divide by SD of all used data from that site), and then the correctly completed trials (hits) were pooled over sessions for further analyses. In total, the analyses used 9 sessions from Monkey K and 14 sessions from Monkey P. All analyses were first calculated per monkey and then combined over monkeys, to give the two animals equal weight. Given that the two monkeys had slightly different gamma and beta peak frequencies, the analyses were aligned to those frequencies (Monkey K: Beta: 18 Hz, Gamma: 74 Hz; Monkey P: Beta: 16 Hz, Gamma: 63 Hz). Following our earlier study ^27^, the analyses were restricted to trials in which the target change occurred first (≈50% of trials), to avoid transients after distractor changes. In addition, to ensure that attention had been fully deployed at the beginning of the analysis window, we excluded trials with target changes less than 800 ms after cue onset.

### Spectral analysis

The main analyses were based on the last 200 ms before target change for power, PPC and GPR, and on the last 400 ms for GC. These signal epochs were Hann tapered, zero padded to 1 s length, and Fourier transformed. Further spectral analyses used the frequency range 1-95 Hz.

The Fourier spectra were squared to obtain the power spectra. The 1/f component of the power spectra was removed using the FOOOF method ^59^.

Phase coherence was quantified as pairwise phase consistency (PPC) ^28^. The PPC was used for site-pair selection. Site pairs whose PPC at >100 Hz exceeded 5SD (of all PPC values of all site pairs of the respective area pair) were considered as affected by artifacts and were excluded; this affected a small percentage of site pairs. Subsequently, site pairs were selected for further analysis, if their PPC spectra at 1-95 Hz exceeded the mean+3SD of all PPC values across all inter-areal site pairs and all frequencies (1-95 Hz).

For each site pair and each trial, the GPR was calculated as the cosine of the deviation of this trial’s phase relation from the mean phase relation of this site pair over all trials.

Directed influences were quantified by calculating frequency-resolved Granger causality (GC)^60^. GC was quantified for all selected site pairs using nonparametric spectral matrix factorization ^61^ of their cross-spectral density matrices, as implemented in the FieldTrip toolbox^58^.

The single-trial correlation between GC and RT was calculated using the jackknife correlation approach ^29^. This approach is based on all possible jackknife replications of trials, that is, all possible leave-one-out subsamples of trials. For each jackknife replication, the average GC and the average RT is calculated. Subsequently, the correlation between those jackknife estimates of GC and RT is calculated. For smooth functions of the data, the jackknife correlation is identical to the regular correlation. For correlations involving GC, the jackknife correlation approach avoids the need to estimate GC for single trials.

### Statistical Analysis

All analyses were performed based on pooled data of both monkeys constituting a fixed-effect analysis that results in inferences on the investigated sample of animals. For this purpose, analysis results were first averaged within each monkey and then averaged over the two monkeys to give the results from each animal equal weight.

All statistical analyses were performed with a multiple-comparison correction based on the Max-based approach ^62^. For each randomization, only the maximal and minimal values across the dimension of multiple comparisons (mostly the frequency dimension) are retained. These values form, after 1000 randomizations, the min-randomization distribution and the max-randomization distribution. The 2.5^th^ percentile of the min-randomization distribution and the 97.5^th^ percentile of the max-randomization distribution were used as significance thresholds. If those thresholds were exceeded by observed, i.e. non-randomized, results, the latter were considered significant with a two-sided false-positive rate of less than 5%, including correction for multiple comparisons.

## ACKNOWLEDGEMENTS

This work was supported by DFG (FOR 1847 FR2557/2-1, FR2557/5-1-CORNET, FR2557/7- 1-DualStreams), EU (HEALTH-F2-2008-200728-BrainSynch, FP7-604102-HBP), a European Young Investigator Award, and the National Institutes of Health (1U54MH091657-WU-Minn-Consortium-HCP). We would like to thank Christini Katsanevaki for help in data preprocessing, and Gustavo Rohenkohl for input on the manuscript.

## DATA AVAILABILITY

Data and code used for analysis in this paper are available from the lead contact on reasonable request.

## LEAD CONTACT

Further information and requests for resources should be directed to and will be fulfilled by the Lead Contact, Pascal Fries.

## AUTHOR CONTRIBUTIONS

M.P., J.V. and P.F. conceptualized the study. C.A.B. and P.F. designed the experiments and performed the implantation surgeries with the help of other colleagues; C.A.B. trained the monkeys and recorded the electrophysiological data; M.P. performed the analyses under supervision of J.V. and P.F.; M.P. and P.F. wrote the paper. All authors reviewed and edited the paper. P.F. supervised and provided funding for this study.

---

**Figure S1.**
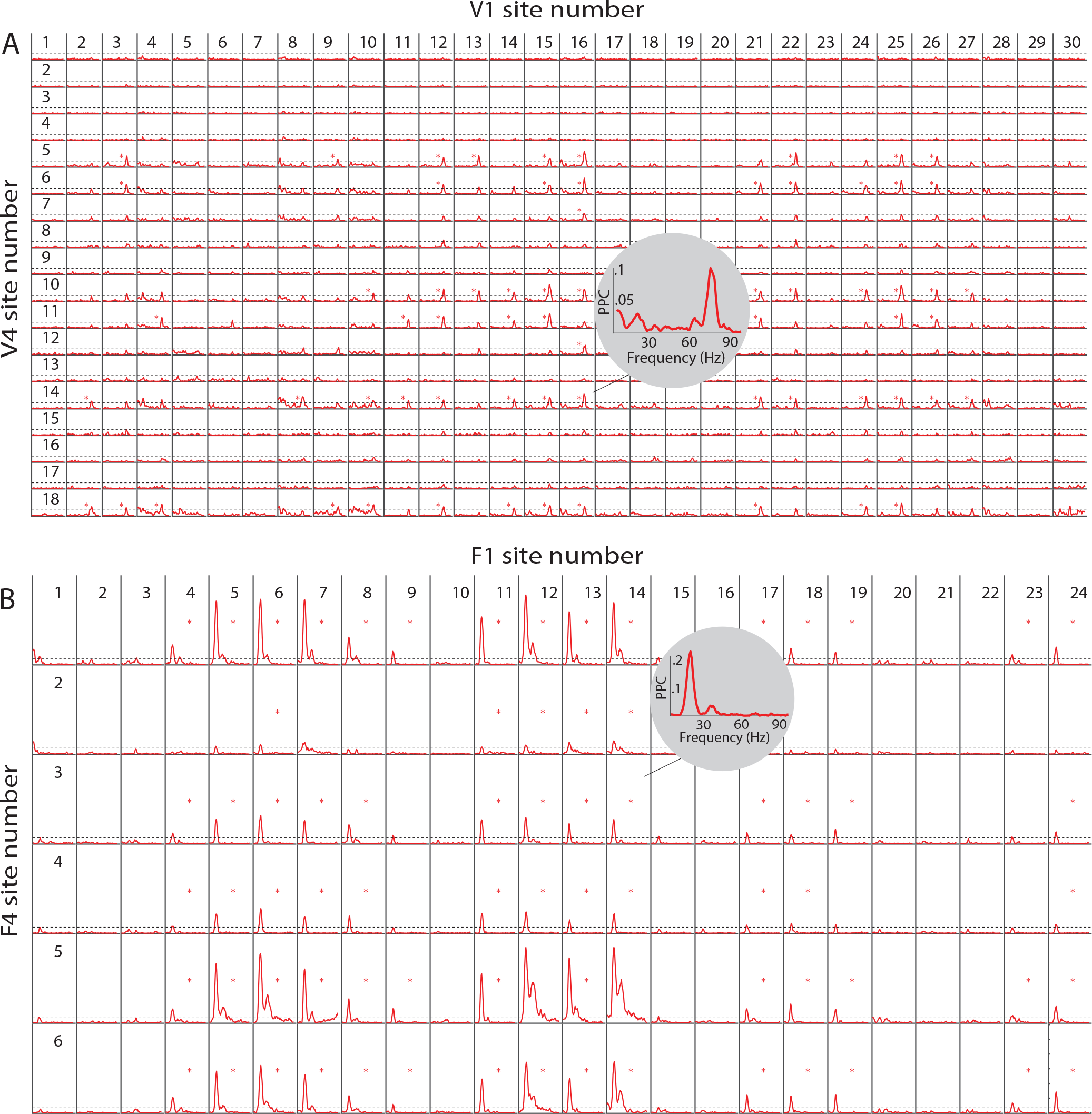
Site-pair selection based on phase coherence. Related to Figures 2 and 3. (A, B) PPC for all interareal site pairs between V1 and V4 (A) or F1 and F4 (B) of monkey K, averaged over conditions IN and OUT. Site pairs with a PPC crossing a threshold (mean+3SD of all PPC values across all frequencies for all site pairs of all recorded areas; dashed lines) were selected for further analyses and are labeled with stars. Insets show zoom-ins for individual selected site pairs from V1-V4 (A) and F1-F4 (B), respectively.

**Figure S2.**
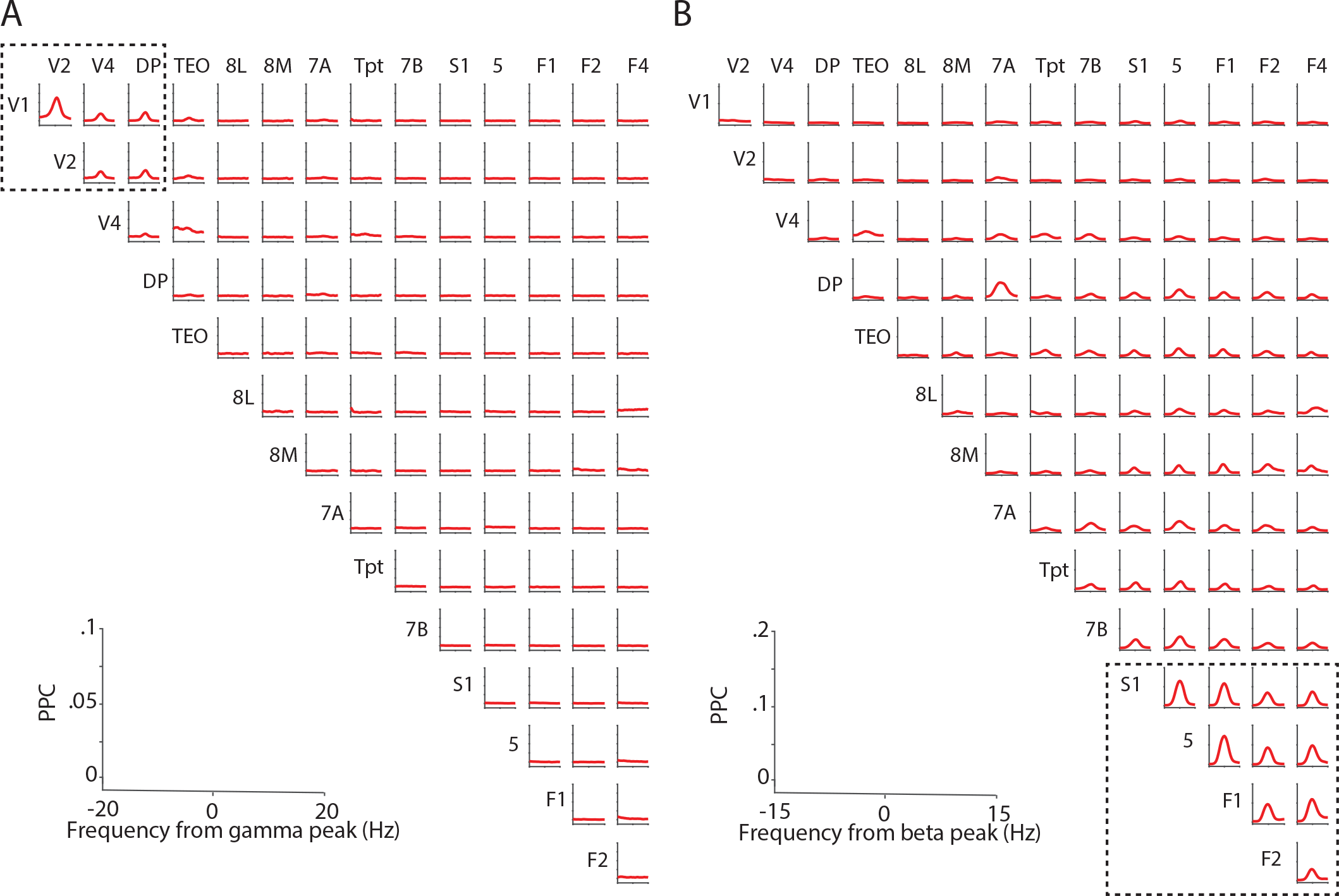
Interareal synchronization in an occipital and fronto-central cluster, in gamma and beta frequencies. Related to Figures 2 and 3. (A, B) PPC averaged over selected site pairs of both monkeys in each area pair, aligned to the individual gamma (A) or beta (B) peak frequency. Areas were ordered according to hierarchical level. We selected area pairs with PPC values exceeding a threshold (mean+2SD of all averaged PPC values across all frequencies and all area pairs). Dashed squares show the occipital gamma cluster (A) and the fronto-central beta cluster (B). The area pair DP-7A showed high beta PPC, but was not a direct neighbor of the beta cluster.

**Figure S3.**
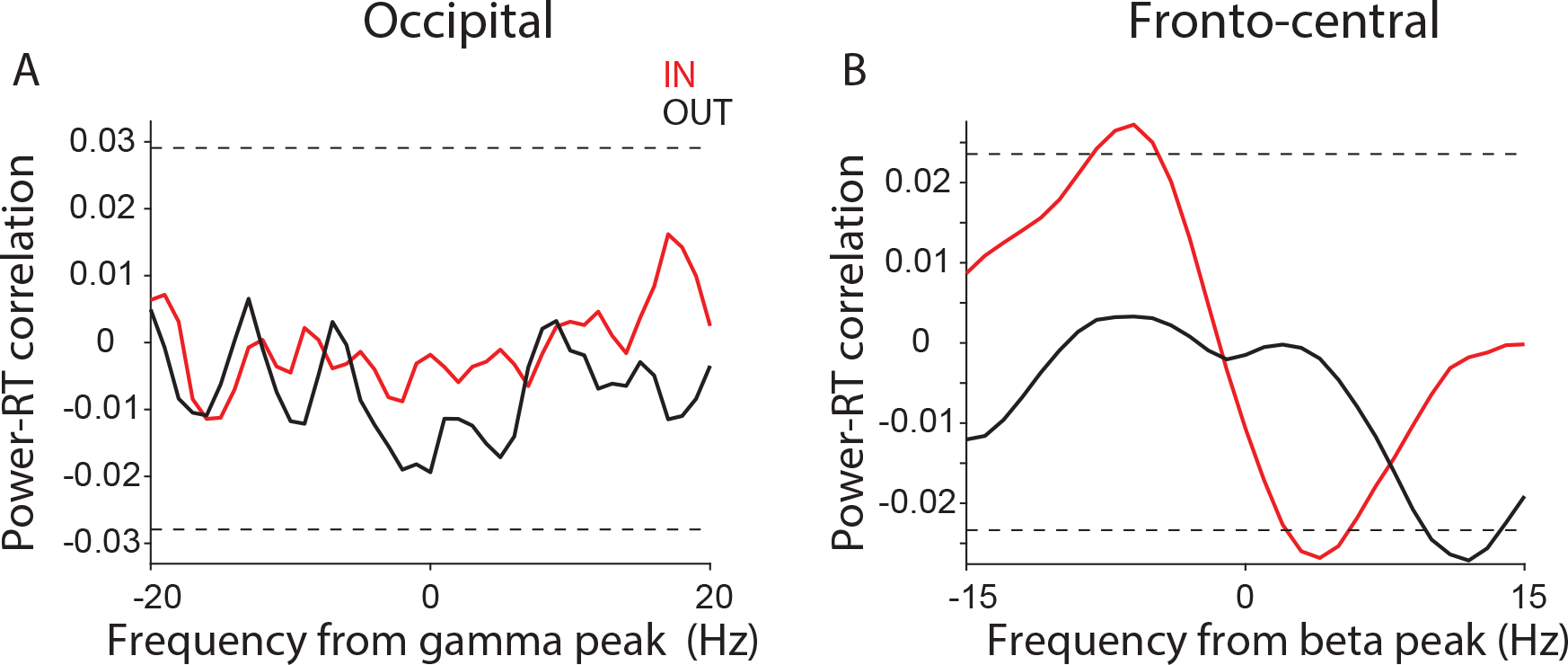
Across-trial correlation between power spectra and RTs. Related to Figure 3. (A, B) Across-trial correlation between power spectra and RTs for IN (red) and OUT (black) conditions, aligned to the gamma peak frequency in the occipital cluster (A) and the beta peak frequency in the fronto-central cluster (B). Correlations were first calculated per site and then averaged over sites.

**Figure S4.**
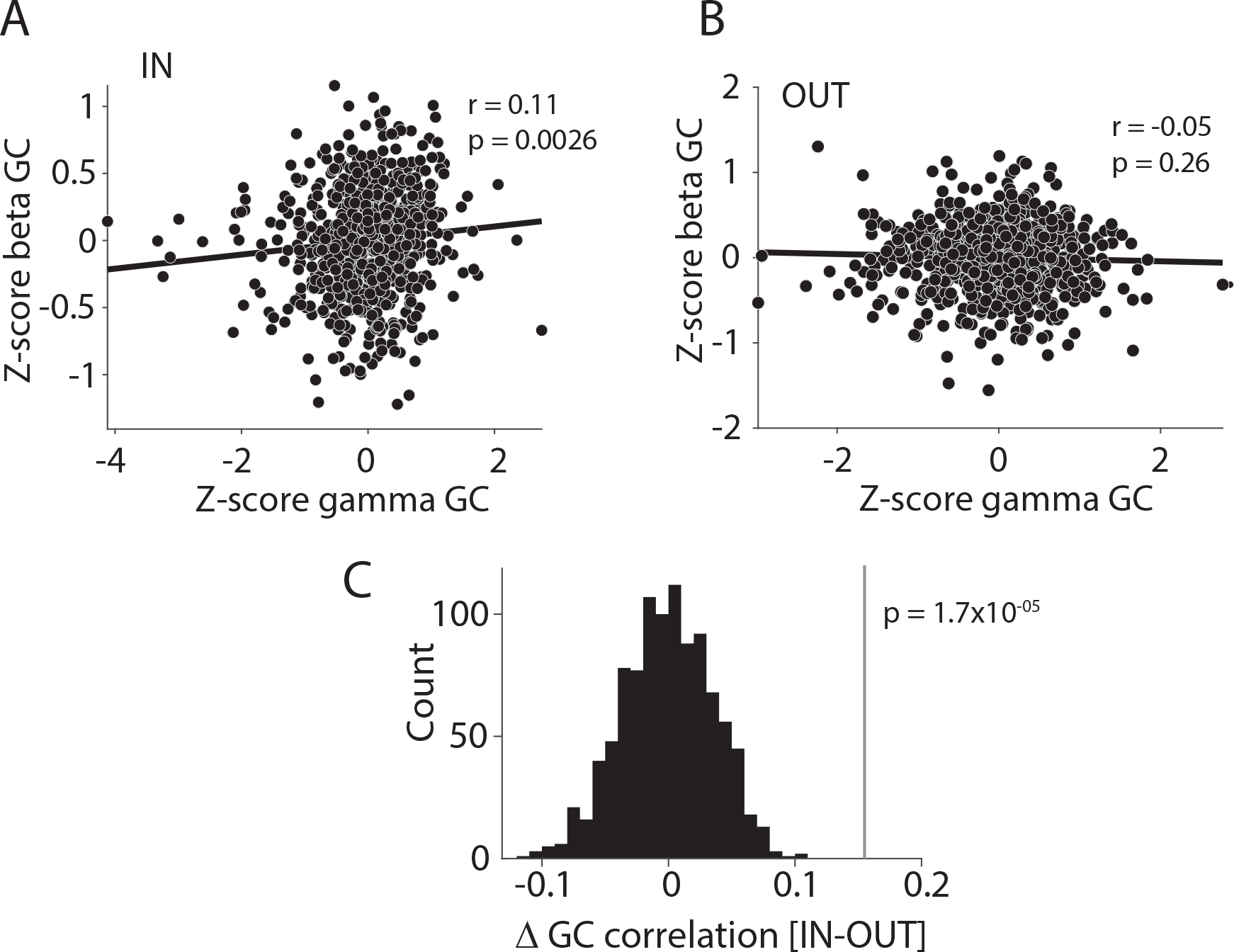
Occipital gamma GC correlates with fronto-central beta GC during IN condition. Related to Figure 5. (A, B) Jackknife correlation (see Methods for details) between occipital gamma GC and fronto-central beta GCs, separately for condition IN (A) and OUT (B). Each dot represents the respective GC values from one jackknife replication, that is, after leaving out one trial, averaged over site pairs of the corresponding areas (gamma GC in occipital areas and beta GC in fronto-central areas). (C) Comparison of the empirically observed difference in GC correlation (IN-OUT) with chance distribution based on 1000 randomizations of trials.

**Figure S5.**
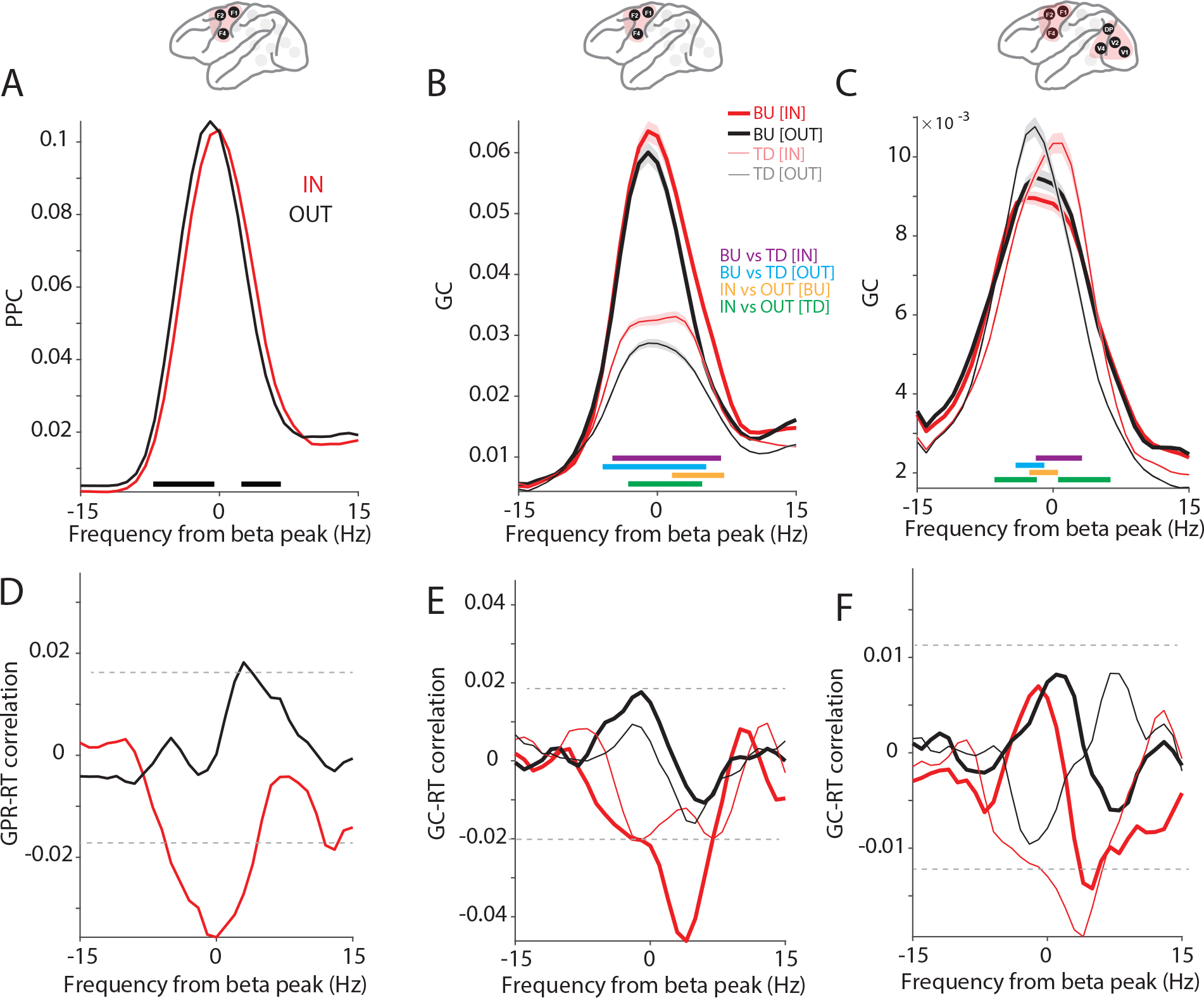
Similar effects for frontal cluster as for fronto-central cluster. (A) Average PPC between area pairs in the frontal cluster (F1, F2, F3; inset), for IN (red) versus OUT (black) conditions. Shaded areas indicate SEM across site pairs. Black horizontal line indicates frequencies with a significant difference between IN and OUT conditions. (B) Interareal GC, averaged over all site pairs of the frontal cluster (inset), aligned to the gamma peak, separately in the bottom-up (BU, tick lines) and top-down (TD, narrow lines) directions, and for the IN (red) and OUT (black) condition. Colored horizontal lines denote significant differences between conditions (IN, OUT, BU, TD), as indicated in the color legend. (C) GC between frontal and occipital clusters aligned to beta peak GC, for IN (red) and OUT (black) conditions, and in bottom-up (BU, tick lines) and top-down (TD, narrow lines) directions, separately. Colored horizontal lines indicate significant frequency bands for the indicated comparisons. (D) Correlation between GPRs and RTs, as a function of frequency for IN (red) and OUT (black) conditions. Black horizontal dashed lines indicate significance thresholds, corrected for multiple comparisons. (E) Jackknife correlation (see Methods for details) between single-trial beta GCs and RTs between the frontal cluster for IN (red) and OUT (black) conditions and in the bottom-up (BU, tick lines) and top-down (TD, narrow lines) directions. (F) Jackknife correlation between single-trial beta GCs and RTs between the occipital and frontal cluster for IN (red) and OUT (black) conditions and in the bottom-up (BU, tick lines) and top-down (TD, narrow lines) directions.

**Figure S6.**
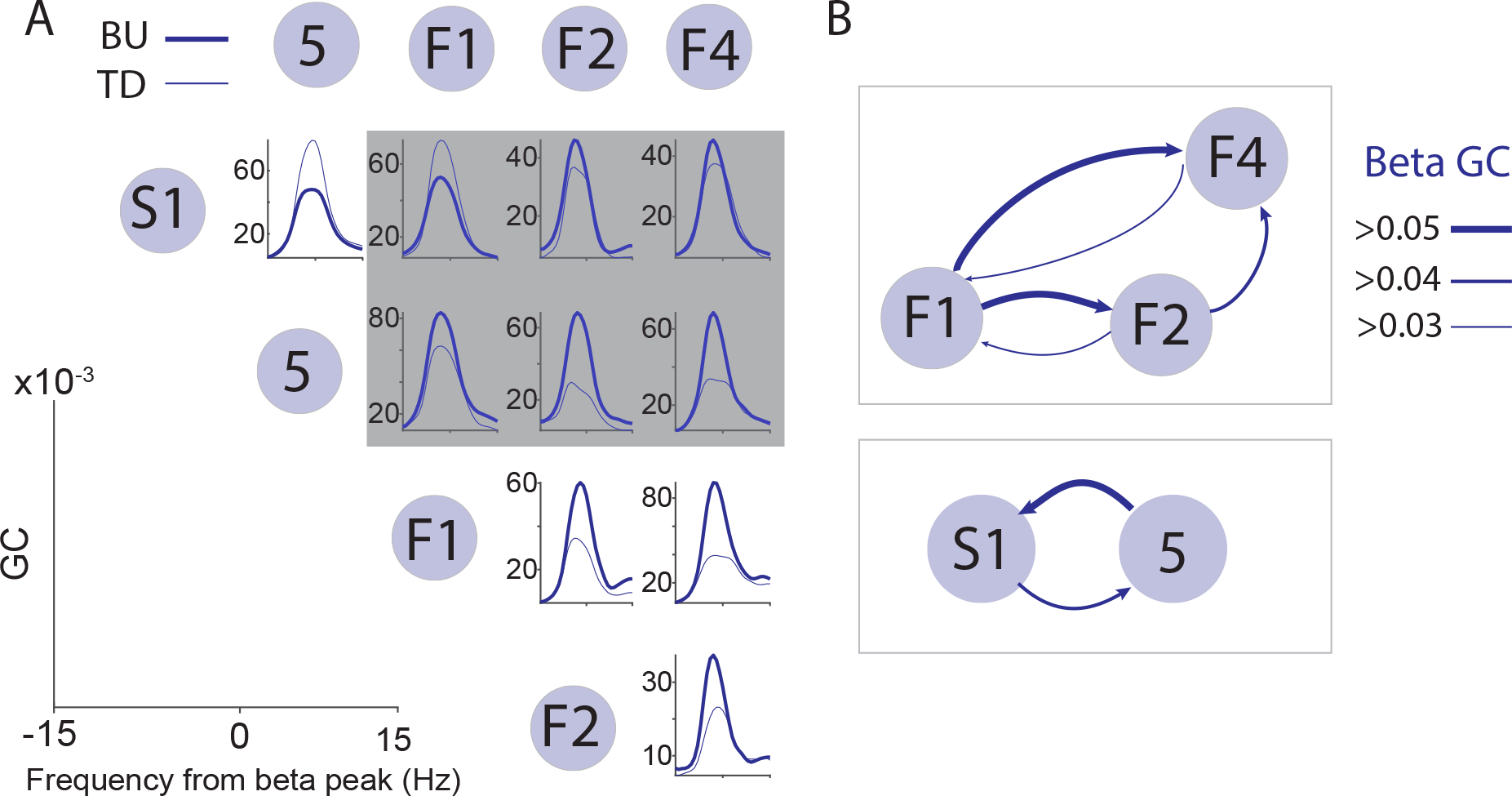
Information flow between fronto-central areas. Related to Figure 7. (A) GC between the area pairs in the fronto-central cluster for bottom-up (BU, tick line) and top-down (TD, narrow line) directions, aligned to the beta peak. Areas were ordered according to hierarchical level. The area pairs highlighted in gray are between areas of different brain systems, namely the somatosensory (S1, 5) and frontal (F1, F2, F4) system. Strength of beta GC between somatosensory and between frontal areas. Between the recorded somatosensory areas, S1 and area 5, beta-band GC was stronger in the top-down direction. Between the recorded frontal areas, F1, F2, F4, beta-band GC was stronger in the bottom-up direction. The strength of GC is indicated by the thickness of the connecting lines.

